# Retrograde transport defects in *Munc18-1* null neurons explain abnormal Golgi morphology

**DOI:** 10.1101/2020.05.12.090811

**Authors:** Annemiek A. van Berkel, Tatiana C. Santos, Hesho Shaweis, Jan R.T. van Weering, Ruud F. Toonen, Matthijs Verhage

## Abstract

Loss of the exocytic Sec1/MUNC18 protein MUNC18-1 or its t-SNARE partners SNAP25 and syntaxin-1 results in rapid, cell-autonomous and unexplained neurodegeneration, which is independent of their known role in synaptic vesicle exocytosis. *cis-*Golgi abnormalities are the earliest cellular phenotypes before degeneration occurs. Here, we investigated whether these Golgi abnormalities cause defects in the constitutive and regulated secretory pathway that may explain neurodegeneration. Electron microscopy confirmed that loss of MUNC18-1 expression results in a smaller *cis-*Golgi. In addition, we now show that *medial*-Golgi and the *trans-*Golgi Network are also affected. However, stacking and cisternae ultrastructure of the Golgi were normal. Overall ultrastructure of null mutant neurons was remarkably normal just hours before cell death occurred. Anterograde ER-to-Golgi and Golgi exit of endogenous and exogenous proteins were normal. In contrast, loss of MUNC18-1 caused reduced retrograde Cholera Toxin transport from the plasma membrane to the Golgi. In addition, MUNC18-1-deficiency resulted in abnormalities in retrograde TrkB trafficking. We conclude that MUNC18-1 deficient neurons have normal anterograde yet reduced retrograde transport to the Golgi. This imbalance in transport routes provides a plausible explanation for the observed Golgi abnormalities and cell death in MUNC18-1 deficient neurons.

**Significance statement:** Loss of MUNC18-1 or its t-SNAREs SNAP25 and syntaxin-1 leads to massive, yet unexplained, neurodegeneration. Previous research showed that Golgi abnormalities are the earliest, shared phenotype. Golgi abnormalities are also an early feature in neurodegenerative diseases, such as Alzheimer’s Disease or Amyotrophic Lateral Sclerosis. This study elucidates the mechanism underlying the Golgi phenotype upon loss of MUNC18-1. By systematically assessing transport routes to and from the Golgi, we show that retrograde endosome-to-Golgi, but not anterograde transport from the Golgi, is disturbed. This imbalance in transport routes provides a plausible explanation for the Golgi phenotype, and may explain the neurodegeneration. The findings in this study contributes to new insights in cellular mechanisms of neurodegeneration.

## Introduction

Synaptic SNARE complex components are known for their essential role in synaptic vesicle fusion in mammalian neurons (Sudhof, 2013). In addition, loss of the target SNAREs (t-SNAREs), SNAP25 and syntaxin-1, or the Sec1/MUNC18 (S/M) protein MUNC18-1 leads to, yet unexplained, neurodegeneration (Verhage et al., 2000; Peng et al., 2013; Vardar et al., 2016; Santos et al., 2017). Their critical role in viability is independent of their role in synaptic vesicle fusion. Viability, but not synaptic transmission, is fully rescued by expression of non-neuronal isoforms SNAP23 in *Snap25* Knock Out (KO) neurons, Syntaxin-2/3/4 in Syntaxin-1 depleted neurons or MUNC18-2/3 in *Munc18-1* KO neurons (Delgado-marti et al., 2007; Peng et al., 2013; Santos et al., 2017). Conversely, several genetic and pharmacological perturbations have been described that block synaptic vesicle fusion and synaptic transmission, but do not cause degeneration (Schiavo et al., 1992; Schoch, 2001; Varoqueaux et al., 2002).

The earliest and shared phenotype among t-SNAREs and Munc18-1 deficient models are abnormalities in Golgi morphology, distinct from apoptosis-induced Golgi abnormalities (Santos et al., 2017). Expression of non-neuronal isoforms does not only rescue viability, but also Golgi morphology, suggesting that these two processes are closely connected (Santos et al., 2017). The Golgi is an essential organelle in the (regulated) secretory pathway. Newly synthesized proteins arrive from the Endoplasmic Reticulum (ER) and move through multiple closely apposed cisternae, allowing posttranslational modifications and subsequent sorting to their final destination (Farquhar and Palade, 1998). This anterograde route is counterbalanced by multiple retrograde trafficking routes to recycle molecules and membrane back to the Golgi (Pavelka and Ellinger, 2008). Golgi structure and function are tightly linked: the vast majority of perturbations in Golgi morphology also affect Golgi function (Chia et al., 2012). A proper equilibrium between anterograde and retrograde routes is essential for cell functioning (Pavelka and Ellinger, 2008).

Here, we tested the hypothesis that the previously observed abnormal Golgi morphology in *Munc18-1* KO neurons leads to Golgi dysfunction and defects in the constitutive and regulated secretory pathway that may explain neurodegeneration. Antero- and retrograde routes in these pathways were studied using electron, live-cell, super-resolution and confocal microscopy. Our data confirm the previously observed *cis-*Golgi abnormalities and extend the phenotype to the Trans Golgi Network (TGN). However, unexpectedly we found that cisternae ultrastructure was normal in *Munc18-1* KO as well as anterograde transport in the secretory pathways. Instead, MUNC18-1 deficiency led to defects in endosome-to-Golgi retrograde pathways. We conclude that loss of Munc18-1 results in disturbances in retro-but not anterograde membrane trafficking pathways. The dysregulation of retrograde trafficking pathways provides a plausible explanation for the previously observed Golgi abnormalities and neurodegeneration.

## Experimental Procedures

### Animals

*Munc18-1* KO mice were generated as described previously (Verhage et al., 2000). On embryonic day 18 (E18) embryos were obtained by caesarean section. All animals were bred and housed according to Institutional and Dutch governmental guidelines.

### Neuronal cultures

Cortices were extracted from E18 embryos and collected in ice-cold Hanks buffered Salt Solution (Sigma) with 7mM HEPES (Invitrogen). After removal of the meninges, neurons were incubated in Hanks-HEPES with 0.25% trypsin (Invitrogen) for 20 min at 37°C. Neurons were washed and triturated with fire polished Pasteur pipettes then counted in a Fuchs-Rosenthal chamber. Neurons were plated in Neurobasal medium supplemented with 2% B-27 (Invitrogen), 1.8% HEPES, 0.25% Glutamax (Invitrogen) and 0.1% Pen/Strep (Invitrogen). Continental (network) cultures were created by plating wild-type (WT) cortical neurons at 25K/well or *Munc18-1* KO neurons at 75K/well. Neurons were plated on 18mm glass coverslips on a layer of rat glia grown on etched glass coverslips applied with 0.1 mg/ml poly-d-lysine and 0.2 mg/ml rat tail collagen (BD Biosciences) solution. For Latrunculin B (LatB) treatment, neurons were plated on a 35mm glass bottom dish. For N-Cadherin immunostaining, neurons were plated on poly-L-ornithine/laminin coated 10mm glass coverslips without glia feeder layer.

### Constructs and lentiviral particles

Constructs encoding pSynapsin-VSVG-EGFP, pSynapsin-NPY-mCherry, pSynapsin-ManII-EGFP (gift Malhotra/Ortega), pSynapsin-MUNC18^WT^-T2A-CreGFP and pSynapsin-MUNC18^V263T^-T2A-CreGFP were subcloned into Lentiviral vectors, and viral particles were produced as described before (Naldini et al., 1996). WT and *Munc18-1* KO neurons were infected with lentiviral particles at 0 days in vitro (DIV). For RUSH experiments, pCMV-Streptavidin / SBP-EGFP-GPI (Addgene #65294, (Boncompain et al., 2012)) and pSynapsin-ManII-ECFP were delivered by standard calcium phosphate precipitation transfection at DIV 1, as described previously (Emperador-Melero et al., 2018).

### Immunocytochemistry

Neuronal cultures were fixed at DIV3 with 3.7% paraformaldehyde (PFA; Electron Microscopy Sciences) then washed 3 times with Phosphate Buffered Saline pH=7.4 (PBS). Neurons were permeabilised with 0.5% Triton X-100, followed by a 30 min incubation in PBS containing 0.1% Triton X-100 and 2% normal goat serum (NGS) to block a-specific binding. All antibodies were diluted in NGS. Neurons were stained with primary antibodies for 2h at room temperature (RT). The following primary antibodies were used: chicken anti-MAP2 (1:1000; Abcam), mouse anti-GM130 (1:1000, Transduction laboratories, rat anti-LAMP1 (1:100, Abcam), rabbit anti-TGN46 (1:500, Abcam), rabbit anti-TrkB (1:500, Millipore), mouse anti-N-Cadherin (1:200, Transduction Laboratory), rabbit anti-mCherry (1:1000, GeneTex), rabbit anti-GFP (1:2000, Bio Connect), rabbit anti-EEA1 (1:50, Cell Signalling). After 3 washes with PBS, neurons were stained with secondary antibodies Alexa Fluor (1:1000; Invitrogen) for 1h at RT. Following 3 additional washes, coverslips were mounted on microscopic slides with Mowiol-DABCO.

For surface staining of TrkB, DIV3 neurons were washed twice in Tyrode’s solution (2 mM CaCl_2_, 2.5 mM KCl, 119 mM NaCl, 2mM MgCl_2_, 20 mM glucose and 25 mM HEPES, pH 7.4) before they were incubated with rabbit anti-TrkB (1:100, Millipore) diluted in Tyrode’s for 15 min at RT. After two washes with Tyrode’s, neurons were fixed in 3.7% PFA for 20 minutes and washed three times with PBS. Neurons were blocked in PBS containing 3% Bovine Serum Albumin (BSA; Acros Organics) for 30 min before incubated with Alexa-647 for 1h at RT. After three washes with PBS, neurons were permeabilized in 0.25% Triton X-100 for 5 min and blocked with 3% BSA for 30 min. Neurons were then stained with primary anti-MAP2 (1:1000) for 2h and secondary Alexa Fluor (1:1000) for 1h. Following 3 additional washes, coverslips were mounted on microscopic slides with Mowiol-DABCO.

For retrograde labelling of Cholera Toxin Subunit B (CTB), neurons were incubated with CTB-488 (100 ng/ml, Thermo Fisher) for 15 minutes at 37°C with 5% CO_2_ in the dark. After washing the cultures with warm Neurobasal, retrograde trafficking was allowed for another 2 hours before neurons were fixed and stained with normal immunocytochemistry protocol.

Images were made on a Zeiss 510 Meta Confocal microscope (Carl Zeiss) with 40X oil immersion objective (NA = 1.3; Carl Zeiss). Z stacks were acquired with 0.5 or 1 μm intervals. Z stacks were collapsed to maximal projections for image analysis.

STED images were made on a Leica SP8 using 100X oil objective (NA = 1.4). Images were deconvoluted using Huygens Professional software. Line segments were drawn in a single focal plane to plot intensity.

### Electron Microscopy

Neurons were fixed for 90 min at RT with 2.5% glutaraldehyde in 0.1M cacodylate buffer, pH 7.4. After washing three times in MilliQ, cells were post-fixed for 90min at RT with 1% OsO_4_/1% KRu(CN)_6_. After dehydration through a series of increasing ethanol concentrations, cells were embedded in Epon and polymerised for 48 h at 60°C. Ultrathin sections (∼80 nm) were cut parallel to the cell monolayer, collected on single-slot, Formvar-coated copper grids, and stained in uranyl acetate and lead citrate in Ultrastainer LEICA EM AC20. Somas were randomly selected at low magnification and imaged using a JEOL1010 transmission electron microscope at 30k to 50k magnification.

### Live-cell imaging

DIV3 neurons were transferred to a NIKON Ti-Eclipse microscope with a Tokai Hit heating system pre-warmed to 37°C with 5% CO_2_. The microscope is equipped with a confocal scanner model A1R+. Time-lapse recordings were made using resonant scanning mode. For the RUSH system, image acquisition was performed every 7 minutes for a total of 120 minutes using a 40X-oil (NA = 1.3) objective. Biotin (40μM, Sigma) was added at t=0. For LatB treatment, imaging was performed every minute for 135 minutes, with a 60X-oil (NA = 1.4) with 2x zoom and z-stacks with 1 μm interval. LatB (20 μM, Sigma) or DMSO (Sigma) as control were added at t=0. In both assays, neurons were imaged in supplemented Neurobasal.

### TrkB antibody uptake assay

The TrkB antibody uptake assay was adapted from (Carrodus et al., 2014). Neurons were incubated with primary rabbit anti-TrkB antibody (1:100; Millipore) diluted in Neurobasal medium at 37°C with 5% CO2 for 1h to promote TrkB internalization. After 2 washes with warm supplemented Neurobasal, neurons were fixed in 3.7% PFA. Cultures were washed in PBS and incubated in 3% Bovine Serum Albumin (BSA) in PBS for 30 min to block a-specific binding. Subsequently, surface TrkB was stained with Alexa-488 (1:1000; Invitrogen) for 1h at RT. After three washing steps in PBS, neurons were incubated for 18h with secondary antibody conjugated with alkaline phosphatase, anti-rabbit-HRP (111-055-003; 1:100; Jackson ImmunoResearch) diluted in 3% BSA to ensure all remaining surface bound anti-TrkB was blocked prior to the permeabilisation steps detecting internalized TrkB fractions. Cultures were washed with PBS and post fixed with 3.7% PFA. Next, neurons were permeabilised with 0.5% Triton X-100 for 5 min and blocked in 3% BSA for 30 min. Internalized TrkB was stained with Alexa-647 (1:1000; Invitrogen), diluted in 3% BSA along with primary chicken anti-MAP2 antibody (1:500; Abcam) for 1.5h at RT. After 3 additional washes, neurons were stained against MAP2 with Alexa-546 (1:1000; Invitrogen) for 1h at RT. Coverslips were mounted on microscopic slides with Mowiol-DABCO and imaged with 40X oil immersion objective (NA = 1.3). Z stacks were acquired with 0.5 μm intervals and collapsed to maximal projections. ImageJ software was used to generate particle masks to analyse puncta area, size and count. The intensity of internalized TrkB was calculated by subtracting surface intensity from the internalized pool using, taking the mean intensity per pixel using ImageJ software.

### Statistical analysis

Statistical analysis and graphing were performed using GraphPad Prism versions 5 and 6 and MATLAB (MathWorks Inc., v2017a). Parametric tests were used whenever assumptions of homoscedasticity and normality were met. Otherwise, non-parametric tests were used. When comparing more than two groups, One-way ANOVA (assumptions met) or Kruskal-Wallis test (assumptions not met) with post-hoc Dunn’s comparisons test was performed. All statistical tests were two-tailed. An error probability level of P<0.05 was accepted as statistically significant. All cellular data is represented in Tukey boxplots. Columns and dots represent individual litters and neurons, respectively.

## Results

### Loss of MUNC18-1 leads to *cis-*Golgi and TGN abnormalities, but normal alignment

Previously, we have shown that loss of MUNC18-1 results in a smaller and rounder *cis-*Golgi (Santos et al., 2017). Here, we confirmed that the GM130 (*cis-*Golgi marker) positive area was ∼33% smaller and ∼50% rounder in DIV 3 *Munc18-1* KO neurons, whereas the mean GM130 intensity was not significantly different (Fig. 1C-E). Staining for TGN46, a TGN marker, revealed a 30% smaller area in *Munc18-1* KO neurons, but the shape of the TGN46 positive area was normal, as quantified with the roundness parameter (Fig. 1F-G).

**Fig 1:**
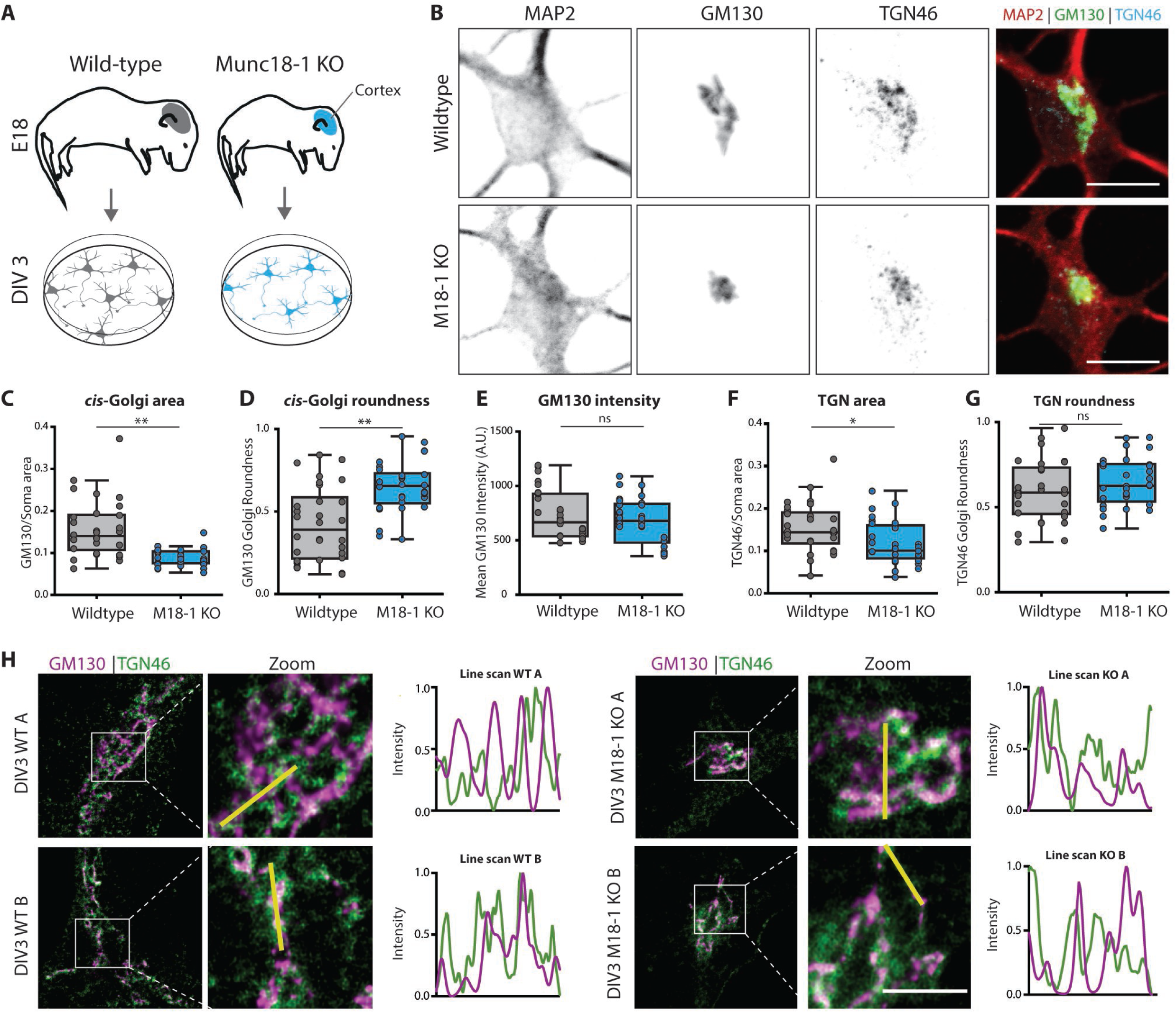
*Munc18-1* KO neurons have a smaller cis and trans Golgi complex. A) Cortical neurons from E18 WT and *Munc18-1* KO mice were plated on glia feeder layers. At DIV3, cultures were fixed and stained for Golgi markers. B) Typical example of WT and *Munc18-1* KO neurons, stained for MAP2 (dendritic marker), GM130 (*cis-*Golgi marker) and TGN46 (TGN marker). Scale bar is 10μm. C) The GM130-positive area was 33% smaller in *Munc18-1* KO neurons compared to WT neurons (p <0.0001, Mann-Whitney U test). D) The GM130-positive area of *Munc18-1* KO neurons was 50% rounder (p<0.0001, unpaired T-test). E) Average intensity of GM130 was comparable between *Munc18-1* KO and WT controls (ns p = 0.27, unpaired T-test). F) *Munc18-1* KO neurons presented a 30% smaller TGN compared to WT neurons (* p = 0.01, unpaired T-test). G) TGN roundness is not affected in *Munc18-1* KO neurons (ns p = 0.22, Mann-Whitney U test). H) Representative STED images and intensity line scans of DIV 3 WT and *Munc18-1* KO neurons, stained for GM130 (*cis-*Golgi marker) and TGN46 (TGN marker). Scale bar is 2μm. Data is represented in Tukey boxplots. Columns and dots represent individual litters and neurons, respectively.

Super-resolution STED imaging was performed to visualize the Golgi abnormalities at higher resolution. Also using STED, GM130 and TGN46-positive areas were smaller in *Munc18-1* KO neurons (Fig. 1H), but the alignment of the GM130 positive areas and TGN46 positive areas was comparable between WT and *Munc18-1* KO. Together, these data confirm *cis*-Golgi abnormalities in *Munc18-1* KO neurons and extend this to the TGN, but indicate a normal spatial alignment of the two compartments.

### Cisternae width and inter-cisterna distance are normal in *Munc18-1* KO neurons

Golgi stack defects, such as dilated cisternae, are frequently observed in mutants that show trafficking defects (Bailey Blackburn et al., 2016; Emperador-Melero et al., 2018). To test for such possible defects, cortical neurons in culture at DIV 1-3 were imaged using electron microscopy (EM) (Fig. 2A). DIV3 is the latest possible time point, since approximately 50% of *Munc18-1* KO neurons has died at this stage and at DIV4 almost all neurons have died (Santos et al., 2017). From DIV 1 onwards, the total Golgi area was smaller in *Munc18-1* KO neurons. At high magnification (50k), individual cisternae had a normal appearance in *Munc18-1* KO neurons (Fig. 2B). Cisternae width and inter-cisternae distance in DIV3 neurons were similar in *Munc18-1* KO and WT neurons (Fig. 2C). The overall morphology and ultrastructure of *Munc18-1* KO neurons were remarkably normal, given the fact that these neurons will die in the next hours (Santos et al., 2017). No signs of neurodegeneration were observed, e.g., membrane blebbing or ruptures, abundant vacuoles or aggregates, or DNA condensation. Ultrastructure of all organelles were indistinguishable from organelles in WT neurons. Together, EM confirms the smaller Golgi area in *Munc18-1* KO neurons, in line with the confocal and STED imaging. However, the overall morphology and ultrastructure, including Golgi cisterna morphology, are normal in *Munc18-1* KO neurons.

**Fig 2:**
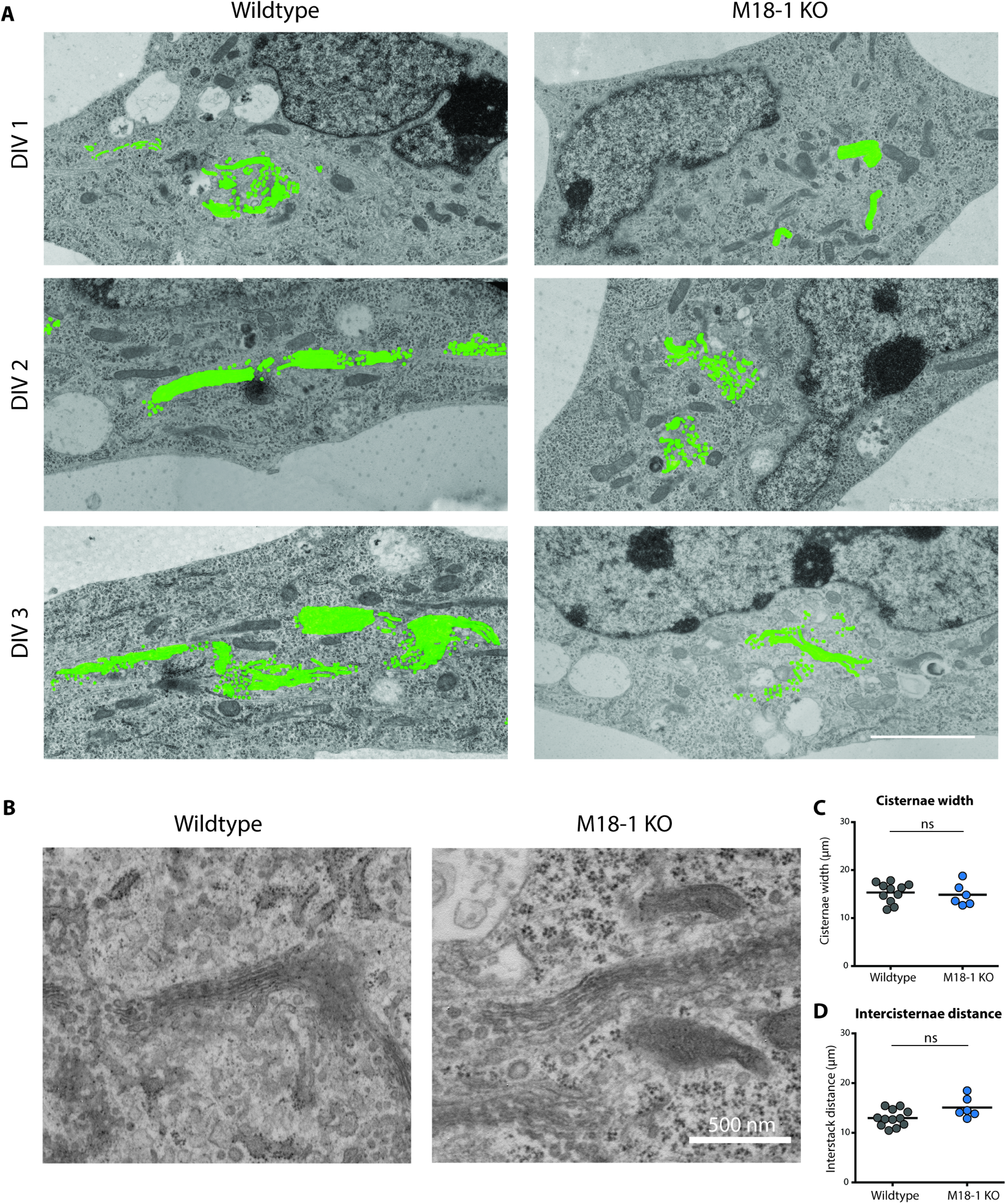
Less Golgi membrane, but normal cisternae structure in *Munc18-1* KO neurons. A) Typical examples of EM images of WT and *Munc18-1* KO neurons fixed at DIV 1, 2 and 3. Golgi membrane is highlighted in green. Scale bar 2.5μm. B) High magnification EM images of Golgi apparatus in WT and *Munc18-1* KO neurons at DIV 3. Scale bar 500 nm. C) Average width of Golgi cisternae was not different between WT and *Munc18-1* KO neurons (ns p = 0.62, Mann-Whitney U test). D) *Munc18-1* KO neurons did not differ in distance between cisternae (ns p = 0.08, Mann-Whitney U test). Mean and individual neurons as dots are shown.

### Actin depolymerisation in WT neurons phenocopies Golgi abnormalities in *Munc18-1* KO neurons

MUNC18-1 is known to regulate F-actin networks (Toonen et al., 2006; Pons-Vizcarra et al., 2019). F-actin is involved in maintaining Golgi morphology (Dippold et al., 2009) and actin depolymerisation by Latrunculin B (LatB) leads to a condensed Golgi (Lázaro-Diéguez et al., 2006), similar to the phenotype observed in *Munc18-1* KO neurons. To test if actin dysregulation causes the *Munc18-1* KO Golgi phenotype, WT and *Munc18-1* KO neurons were treated with LatB (20µM). ManII-GFP was used as a live marker for the medial-Golgi. In *Munc18-1* KO neurons, the ManII-GFP positive area was ∼65% smaller, as expected (Fig. 3B-C). LatB treatment in WT neurons resulted in a gradual but marked reduction in the ManII-GFP positive area, cumulating in a ∼40% total reduction 2h after LatB addition. However, in *Munc18-1* KO neurons, LatB had no effect on the ManII-GFP positive area. Hence, actin depolymerisation in WT neurons phenocopies the *Munc18-1* KO Golgi phenotype, but does not further reduce Golgi area in *Munc18-1* KO neurons, suggesting that dysregulation of F-actin may indeed contribute to Golgi abnormalities in *Munc18-1* KO neurons.

**Fig 3:**
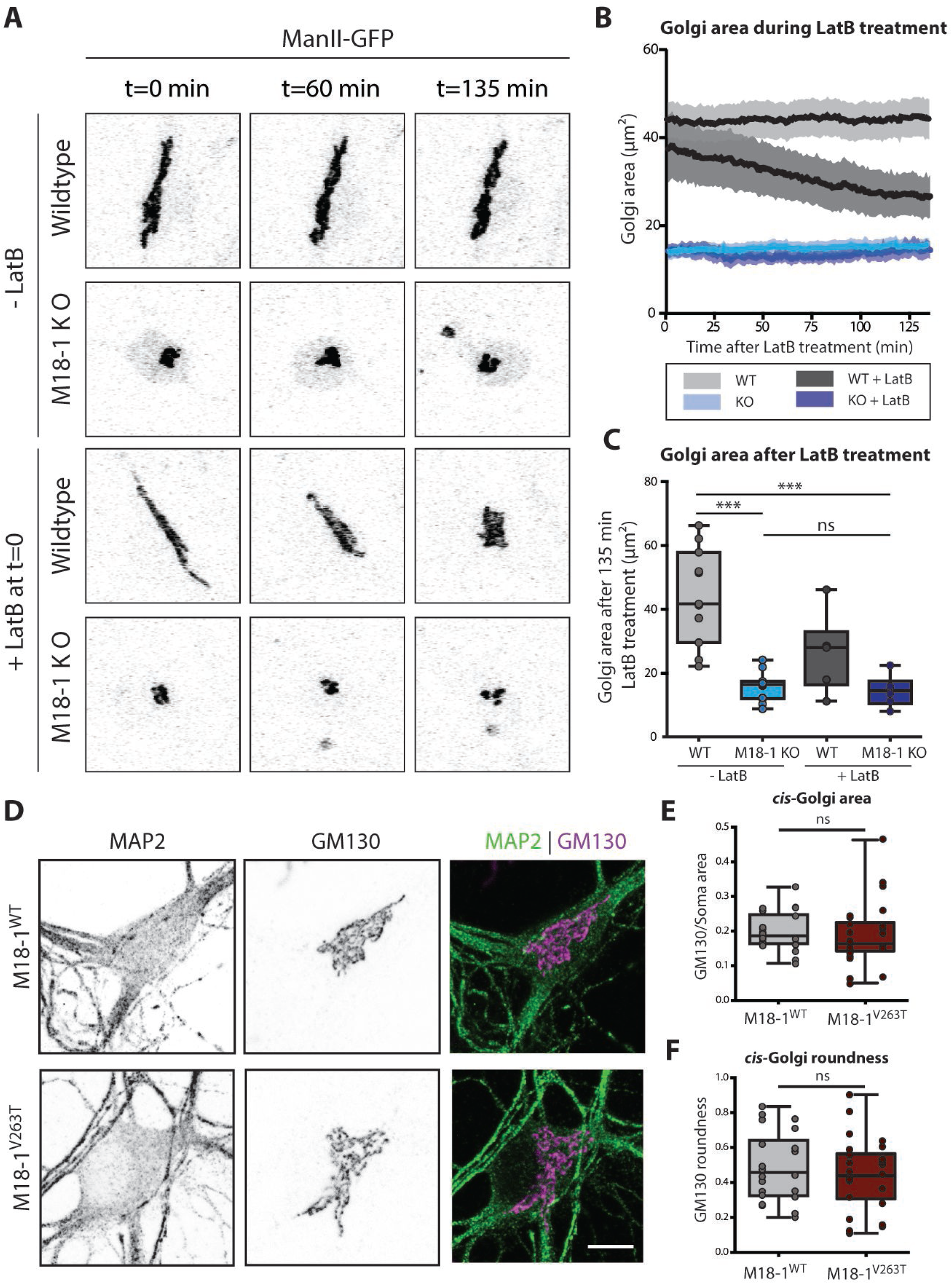
Actin depolymerisation reduces Golgi area in WT but not in *Munc18-1* KO neurons. A) WT and *Munc18-1* KO neurons were infected with ManII-GFP (Golgi marker) and treated with LatB at DIV3. Golgi morphology was imaged for 135 minutes. B) Golgi area in WT and *Munc18-1* KO neurons with and without LatB treatment over time. C) Average Golgi area in WT and *Munc18-1* KO neurons with and without LatB treatment after 135 minutes of treatment. WT Golgi area was significantly larger than *Munc18-1* KO with and without LatB treatment (p<0.0001, Kruskal-Wallis test with post-hoc Dunn’s comparisons test). Golgi area of WT + LatB treatment was similar as in *Munc18-1* KO neurons (p>0.05, Kruskal-Wallis test with post-hoc Dunn’s comparisons test). In *Munc18-1* KO neurons, LatB did not decrease Golgi area (p>0.05, Kruskal-Wallis test with post-hoc Dunn’s comparisons test). D) Typical examples of *Munc18-1* KO neurons rescued with M18-1^WT^ and M18-1^V263T^ at DIV14 stained for MAP2 (dendritic marker) and GM130 (*cis-*Golgi marker). Scale bar is 10μm. E) The GM130-positive area in M18-1^V263T^ was not different from M18-1^WT^ neurons (ns p=0.27, Mann-Whitney U test). F) GM130-positive area roundness was not affected in M18-1^V263T^ neurons (ns p=0.38, unpaired T-test). Data is represented in Tukey boxplots. Columns and dots represent individual litters and neurons, respectively.

To test this possibility further, a MUNC18-1 mutant (V263T) was expressed in *Munc18-1* KO neurons known to rescue secretion and Syntaxin-1 targeting, but not F-actin dysregulation (Pons-Vizcarra et al., 2019). *Munc18-1* KO neurons were infected with MUNC18-1^WT^ or MUNC18-1^V263T^ and *cis-*Golgi morphology was examined at DIV14 (Fig. 3D). MUNC18-1^V263T^ neurons showed normal survival. GM130-positive area and roundness were not affected by the V263T mutation (Fig. 3E-F). Together, these data show that MUNC18-1’s role in F-actin regulation is not causally related to its function in maintaining Golgi morphology.

### Endogenous proteins of the secretory pathway do not accumulate in the Golgi of *Munc18-1* KO neurons

To understand whether the abnormal Golgi morphology affects Golgi function, we examined the expression levels and localisation of two proteins that are trafficked through the Golgi: cell-adhesion protein N-Cadherin and Neurotrophin receptor Tropomyosin Receptor Kinase B (TrkB) (Fig4. A-B) (Klein et al., 1991; Wiertz et al., 2011). In *Munc18-1* KO neurons, the N-Cadherin staining intensity in the TGN46-positive area was comparable to WT neurons (Fig. 4C), while the TrkB intensity was ∼55% lower (Fig. 4D). However, these differences between N-Cadherin and TrkB staining were not specific for the Golgi marker areas. The total staining intensity in the dendrites and soma was also similar for N-Cadherin, but also ∼55% reduced for TrkB (Fig. 4E-F). Hence, TrkB levels are reduced in *Munc18-1* KO neurons, not specifically in the Golgi and no evidence for specific Golgi retention was obtained for either marker.

**Fig 4:**
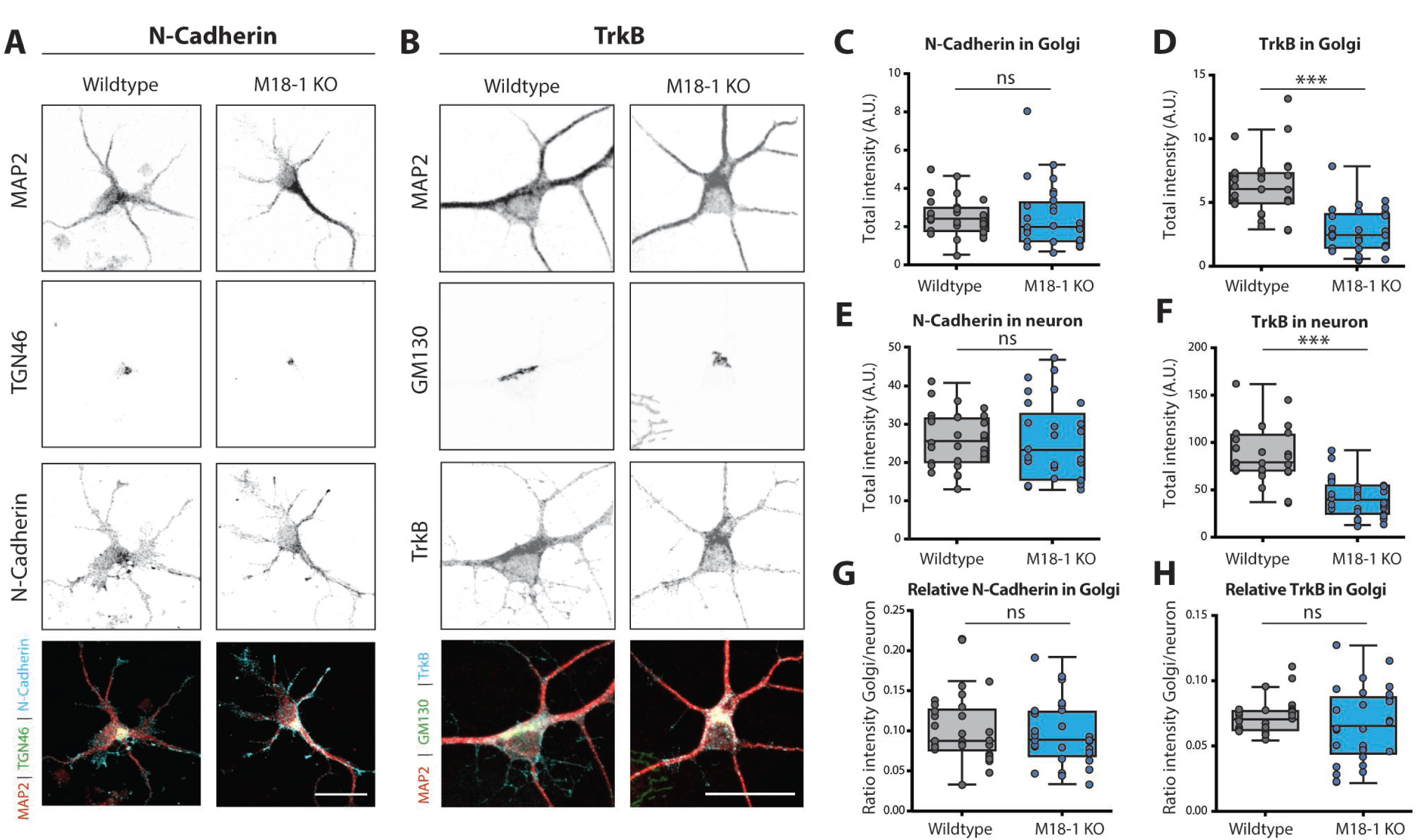
Endogenous markers of the secretory pathway do not accumulate in the Golgi of *Munc18-1* KO neurons. A) Typical example of WT and *Munc18-1* KO neurons at DIV3 stained for MAP2 (dendritic marker), TGN46 (Trans Golgi Network Marker) and N-Cadherin. Scale bar is 25μm. B) Typical example of WT and *Munc18-1* KO neurons at DIV3 stained for MAP2 (dendritic marker), GM130 (*cis-*Golgi marker) and TrkB. Scale bar is 25μm. C) N-Cadherin intensity in Golgi was not different between WT and *Munc18-1* KO (ns p = 0.24, Mann-Whitney test). D) TrkB intensity in Golgi was ∼55% lower in *Munc18-1* KO (p < 0.0001, Mann-Whitney test). E) *Munc18-1* KO neurons did not differ in total N-Cadherin intensity (ns p = 0.82, unpaired T-test). F) Total TrkB intensity in *Munc18-1* KO neurons was ∼55% lower (p < 0.0001, unpaired T-test). G) Relative N-Cadherin distribution in the Golgi, measured by N-Cadherin intensity in Golgi (C) divided by total N-Cadherin intensity (E), was not different in *Munc18-1* KO neurons (ns p = 0.65, Mann-Whitney test). H) Relative TrkB distribution in the Golgi, measured by TrkB intensity in Golgi (D) divided by total TrkB intensity (F), was not different in *Munc18-1* KO neurons (ns p = 0.36, Mann-Whitney test). Data is represented in Tukey boxplots. Columns and dots represent individual litters and neurons, respectively.

To confirm this conclusion quantitatively, TrkB staining intensity in GM130-positive area was divided by the staining intensity in the total neuron. For both N-Cadherin and TrkB, no differences were found in the relative levels in the GM130-positive areas compared to the rest of the neuron (Fig. 4G-H). Together, these data show that although some proteins show lower expression levels, no evidence was obtained for specific Golgi retention, suggesting that Golgi function is not affected by its smaller size and altered shape in *Munc18-1* KO neurons.

### ER-to-Golgi and Golgi exit cargo trafficking is not affected upon MUNC18-1 loss

To examine whether specific secretory pathways are disturbed in *Munc18-1* KO neurons, we overexpressed cargo proteins for the constitutive (VSVG-GFP) and regulated (NPY-mCherry) pathway (Horton and Ehlers, 2003; Persoon et al., 2018). Expression levels of VSVG-GFP and NPY-mCherry were 60%-55% lower in the GM130-positive area of *Munc18-1* KO neurons, respectively (Fig. 5C-D). Quantification of expression in dendrites and soma showed a similar reduction in VSVG-GFP (∼60%, Fig. 5E) and NPY-mCherry levels (∼45%, Fig. 5F) in *Munc18-1* KO neurons. Hence, similar to the situation for endogenous cargo (Fig. 4), the reduced expression of heterologous cargo in the Golgi area appears to be a consequence of overall lower expression levels, rather than Golgi-retention.

**Fig 5:**
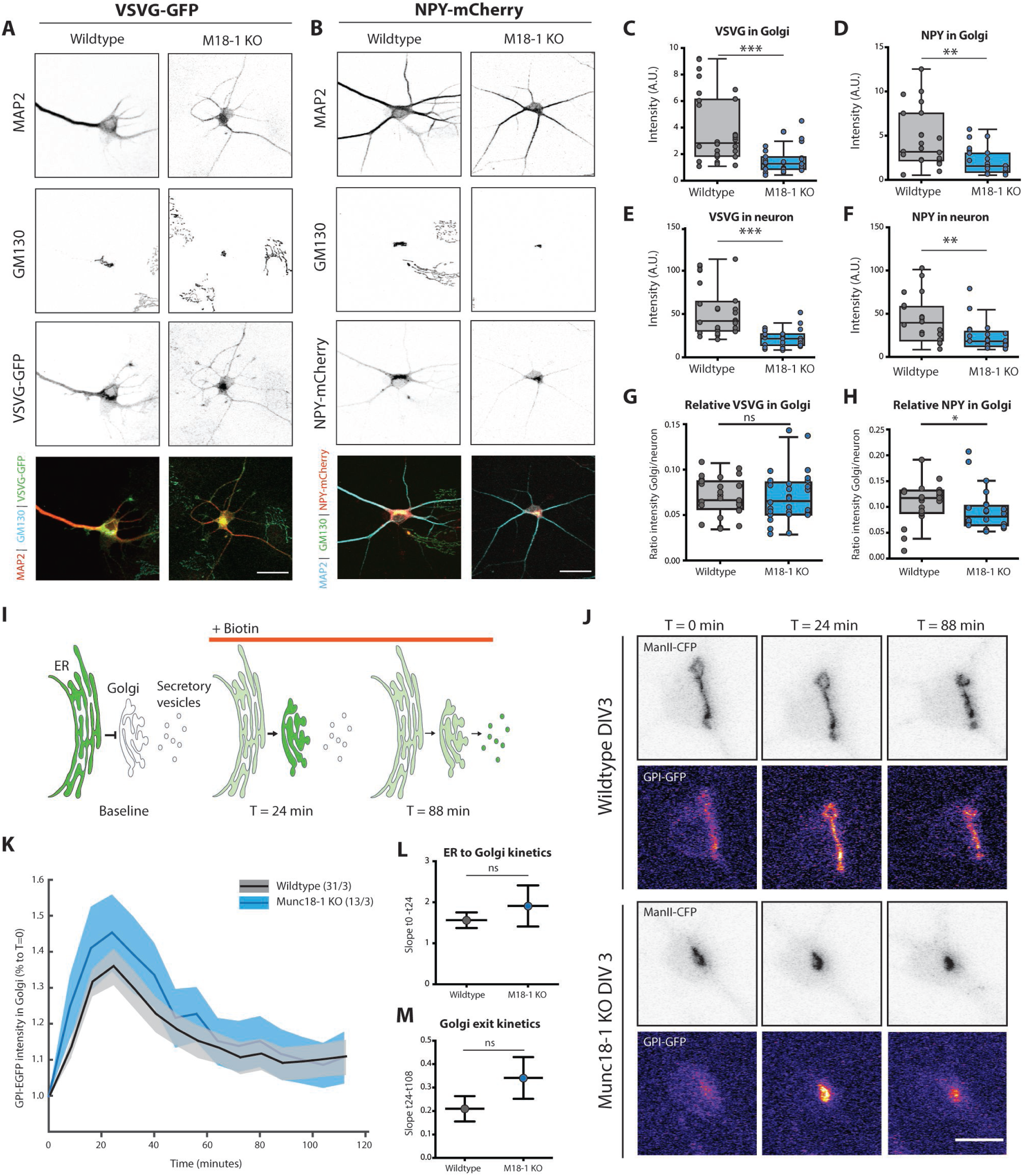
Anterograde transport through the Golgi is unaffected in *Munc18-1* KO neurons. A) Typical example of WT and *Munc18-1* KO neurons overexpressing VSVG-GFP at DIV3, stained for MAP2 (dendritic marker), *cis-*Golgi (GM130) and GFP. Scale bar is 25μm. B) Typical example of WT and *Munc18-1* KO neurons overexpressing NPY-mCherry at DIV 3, stained for MAP2 (dendritic marker), *cis-*Golgi (GM130) and mCherry. Scale bar is 25μm. C) VSVG-GFP intensity in the Golgi of *Munc18-1* KO neurons was ∼60% lower (p < 0.0001, Mann-Whitney test) D) *Munc18-1* KO neurons showed a ∼55% reduction in NPY-mCherry intensity in the Golgi area (p = 0.001, Mann-Whitney test). E) VSVG-GFP intensity in neuron was ∼60% lower in *Munc18-1* KO neurons (p<0.0001, Mann-Whitney test). F) NPY-mCherry intensity in *Munc18-1* KO neurons was ∼45% lower (p = 0.0024, Mann-Whitney test). G) Relative VSVG-GFP distribution in the Golgi, measured by VSVG-GFP intensity in Golgi (C) divided by total VSVG-GFP intensity (E), was not different in *Munc18-1* KO neurons (ns p = 0.58, Mann-Whitney test). H) Relative NPY-mCherry distribution in the Golgi, measured by NPY-mCherry intensity in Golgi (D) divided by total NPY-mCherry intensity (F), was 15% lower in *Munc18-1* KO neurons (ns p = 0.026, Mann-Whitney test). Data is represented in Tukey boxplots. Columns and dots represent individual litters and neurons, respectively. I) Cartoon representing RUSH assay. Under baseline conditions, GPI-GFP-SBP is bound to KDEL-streptavidin and therefore retained in the ER. Upon biotin administration, the binding between SBP and streptavidin is reversed, thereby releasing the GPI-GFP cargo into the secretory pathway. J) Representative images of WT and *Munc18-1* KO DIV3 neurons expressing Golgi marker ManII-CFP and GPI-GFP. Biotin was added at T = 0. Scalebar is 10μm. K) GPI-GFP intensity in the Golgi of WT and *Munc18-1* KO neurons over time after biotin administration. Shown are mean and SEM. WT = 31 neurons/3 independent litters. KO = 13 neurons/3 independent litters. L) GPI transport to the Golgi, measured as the fitted slope between t=0 and the highest intensity, t=24min, was not significantly different between WT and *Munc18-1* KO neurons (p=0.42, F=0.633, DFn=1, DFd=172, linear regression). M) No differences were observed in Golgi exit, measured as the fitted slope between t=24 and t=112, between the two conditions (p=0.19, F=1.656, DFn=1, DFd=480, linear regression).

To confirm this conclusion again quantitatively, relative expression levels overlapping with GM130 staining, were divided by the average fluorescence intensity in the total neuron. For VSVG-GFP, the relative levels in the Golgi were not different in *Munc18-1* KO neurons (Fig. 5G) and ∼15% lower for NPY-mCherry (Fig. 5H). Hence, overall expression levels of markers for the constitutive and regulated secretory pathway are lower in *Munc18-1* KO neurons, but no evidence was observed for Golgi retention of either marker.

The RUSH system was used to examine cargo trafficking in live neurons (Boncompain et al., 2012). GPI-GFP, a marker of the constitutive secretory pathway, fused to Streptavidin binding protein (SBP) and the ER target peptide sequence KDEL fused to Streptavidin were expressed in *Munc18-1* KO and WT neurons. Under baseline conditions, SBP and Streptavidin bind, trapping GPI-GFP in the ER. Biotin administration reverses the binding, allowing GPI-GFP to travel through the secretory pathway (Fig. 5I-J). Before adding biotin, GPI-GFP was similarly localised in WT and *Munc18-1* KO neurons (Fig. 5J, T = 0 min). Addition of biotin resulted in rapid translocation of GPI-GFP to ManII-CFP positive Golgi areas, reaching a maximum after 24 minutes (Fig. 5K). No differences in the kinetics of ER to Golgi transport were detected between WT and *Munc18-1* KO neurons (Fig. 5K-L). Subsequent Golgi exit resulted in decreasing GPI-GFP intensity in the ManII-CFP areas. *Munc18-1* KO neurons performed equally well in Golgi exit as WT neurons (Fig. 5K-M). Taken together, these data indicate that the altered Golgi size and morphology in *Munc18-1* KO neurons does not affect ER-to-Golgi transport and Golgi exit, suggesting that anterograde trafficking in the two main secretory routes is normal in the absence of Munc18-1.

### Retrograde Golgi pathways are affected in *Munc18-1* KO neurons

Retrograde trafficking pathways from the endolysosomal system to the Golgi are important to replenish essential proteins and lipids to the Golgi, which is essential for survival (Bonifacino and Rojas, 2006). Impaired retrograde transport may explain the Golgi abnormalities and cell death in *Munc18-1* KO neurons. First, we examined the morphology of the endolysosomal pathway. LAMP1 is an integral component of lysosomes, and often used as a marker for lysosomes (Fig. 6A) (Chen et al., 1988). LAMP1-positive area and intensity in the soma were unaffected in *Munc18-1* KO neurons (Fig. 6B-C). Staining for early endosome antigen 1 (EEA1), a widely used marker for early endosomes, showed a punctate pattern in soma and dendrites (Fig. 6D). MUNC18-1 depletion did not affect the number of EEA1 puncta nor the size of EEA1-positive puncta (Fig. 6E-F). These data indicate that, unlike the Golgi, endolysosomal morphology is normal upon deletion of MUNC18-1 expression.

**Fig 6:**
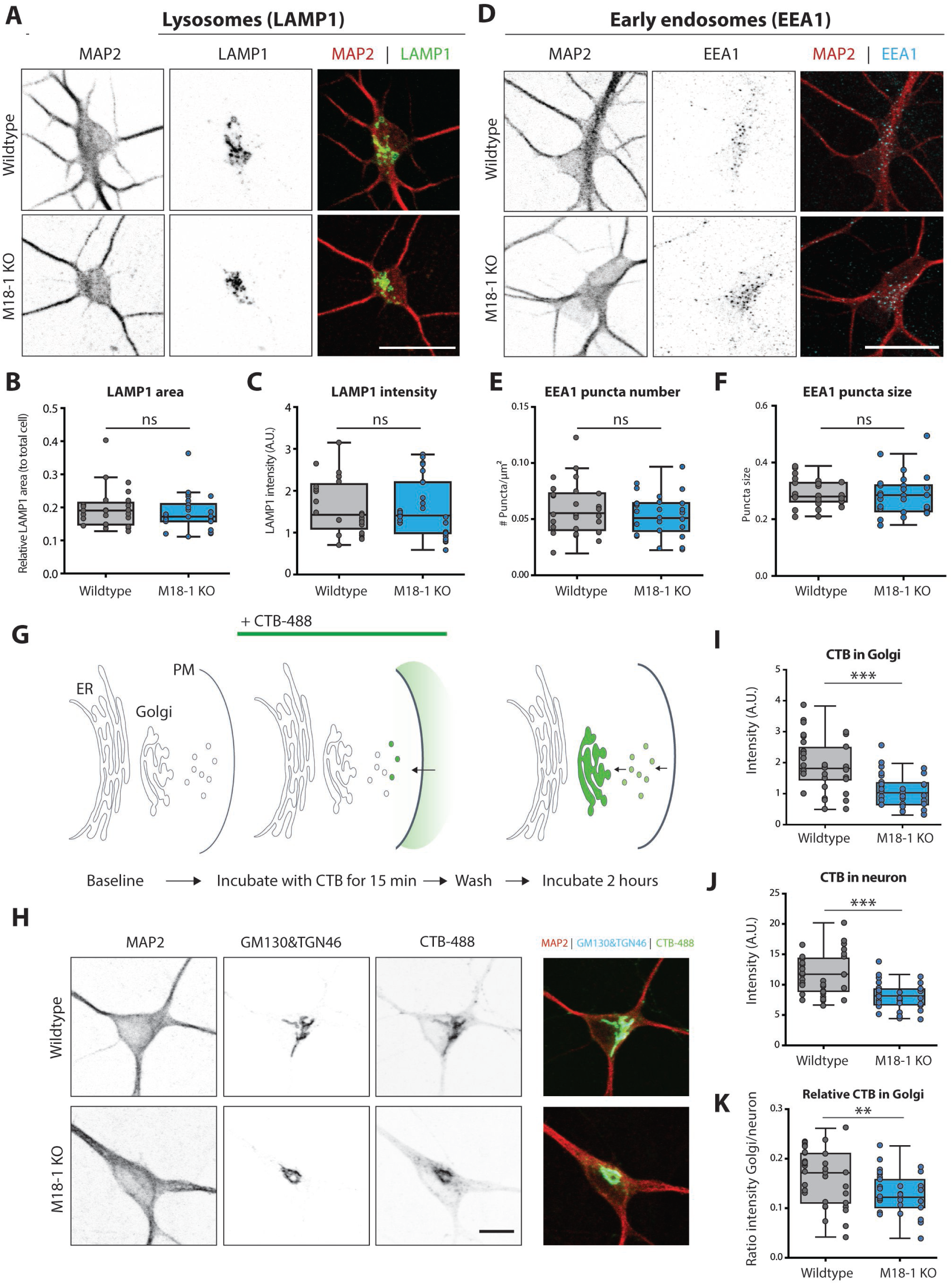
*Munc18-1* KO neurons show reduced endosome-to-Golgi retrograde transport of Cholera Toxin B. A) Typical example of WT and *Munc18-1* KO neurons at DIV3 stained for MAP2 (dendritic marker) and LAMP1 (lysosomes). Scale bar is 25μm. B) The relative LAMP1 area, measured as the LAMP1-positive area over soma area, was not changed in *Munc18-1* KO neurons (ns p=0.41, Mann-Whitney test). C) No differences were observed in LAMP1 intensity between both conditions (ns p=0.81, unpaired T-test). D) Typical example of WT and *Munc18-1* KO neurons at DIV 3 stained for MAP2 (dendritic marker) and EAA1 (early endosomes). Scale bar is 25μm. E) Number of EEA1-positive puncta per μm^2^ was unchanged in *Munc18-1* KO neurons (ns p=0.34, unpaired T-test). F) The size of EEA1-positive puncta was comparable between the two conditions (ns p=0.42, Mann-Whitney test). G) Cartoon representing the CTB assay. After a 15-minute incubation of CTB, uninternalized CTB is washed out. Internalized CTB is allowed to retrogradely traffic to the Golgi for 2 hours. H) Typical examples of WT and *Munc18-1* KO neurons fixed after the CTB assay. Neurons were stained for MAP2 (dendritic marker) and GM130&TGN46 (*cis-*Golgi and TGN markers). I) CTB intensity in Golgi was ∼45% lower in *Munc18-1* KO neurons (p < 0.0001, Mann-Whitney test). J) Total CTB internalized in *Munc18-1* KO neurons was ∼30% lower (p < 0.0001, Mann-Whitney test). K) Relative CTB distribution in the Golgi, measured by CTB intensity in Golgi (I) divided by total CTB intensity (J), was 20% lower in *Munc18-1* KO neurons (p = 0.004, unpaired T-test). Data is represented in Tukey boxplots. Columns and dots represent individual litters and neurons, respectively.

Cholera Toxin Subunit B (CTB) fused to Alexa-488 was used to investigate retrograde transport (Fig. 6G). CTB travels from the plasma membrane (PM) to early endosomes, recycling endosomes and the Golgi (Wernick et al., 2010; Matsudaira et al., 2015; Wang et al., 2016). After two hours of CTB incubation, the majority of the CTB was translocated from the PM to GM130/TGN46-positive areas in the soma (Fig. 6H). CTB intensity in these Golgi areas was ∼45% lower in *Munc18-1* KO neurons than in WT neurons (Fig. 6I). Total CTB-Alexa-488 internalized in *Munc18-1* KO neurons showed a ∼30% reduction (Fig. 6J). The relative distribution of CTB in the GM130/TGN46 positive areas, as calculated by the ratio of CTB-Alexa-488 fluorescence in GM130/TGN46-positive areas over total CTB fluorescence, was ∼20% lower in *Munc18-1* KO neurons (Fig. 6K). Hence, retrograde transport of CTB to the Golgi is impaired in *Munc18-1* KO neurons.

### Abnormal retrograde TrkB trafficking in *Munc18-1* KO neurons

To further investigate the impaired retrograde trafficking, we examined the retrograde endosomal trafficking of TrkB using live labelling with a TrkB antibody (Carrodus et al., 2014; Barford et al., 2017; Budzinska et al., 2020). Before antibody incubation, the mean TrkB surface labelling was similar in WT and *Munc18-1* KO neurons (Fig. 7A-B). Subsequently, live neurons were incubated with a TrkB antibody. To distinguish surface from internalized TrkB, both fractions were stained with a different secondary antibody (Fig. 7C). In both WT and *Munc18-1* KO neurons a punctate pattern of internalized TrkB was observed (Fig. 7D). The mean TrkB punctum intensity in *Munc18-1* KO neurons was ∼30% and ∼35% lower in the soma and dendrites, respectively (Fig. 7E-F). Conversely, the number of TrkB puncta was ∼20% higher in the soma of *Munc18-1* KO neurons, but not different in neurites (Fig. 7G-H). Puncta size was increased by 15% and 10% in soma and dendrites, respectively (Fig. 7I-J). The total pool of internalized TrkB, measured by multiplying punctum intensity by total puncta area, remained unaffected in *Munc18-1* KO neurons (Fig. 7K). These data together show that endocytosed TrkB puncta were bigger and contained less TrkB in *Munc18-1* KO neurons. Moreover, the density of TrkB puncta was higher in the soma, but not in dendrites, while the total pool of internalized TrkB was unaffected. Together with the impaired retrograde CTB transport, we conclude that retrograde trafficking is affected by MUNC18-1 deficiency and may explain the morphological abnormalities in the Golgi in *Munc18-1* KO neurons.

**Fig 7:**
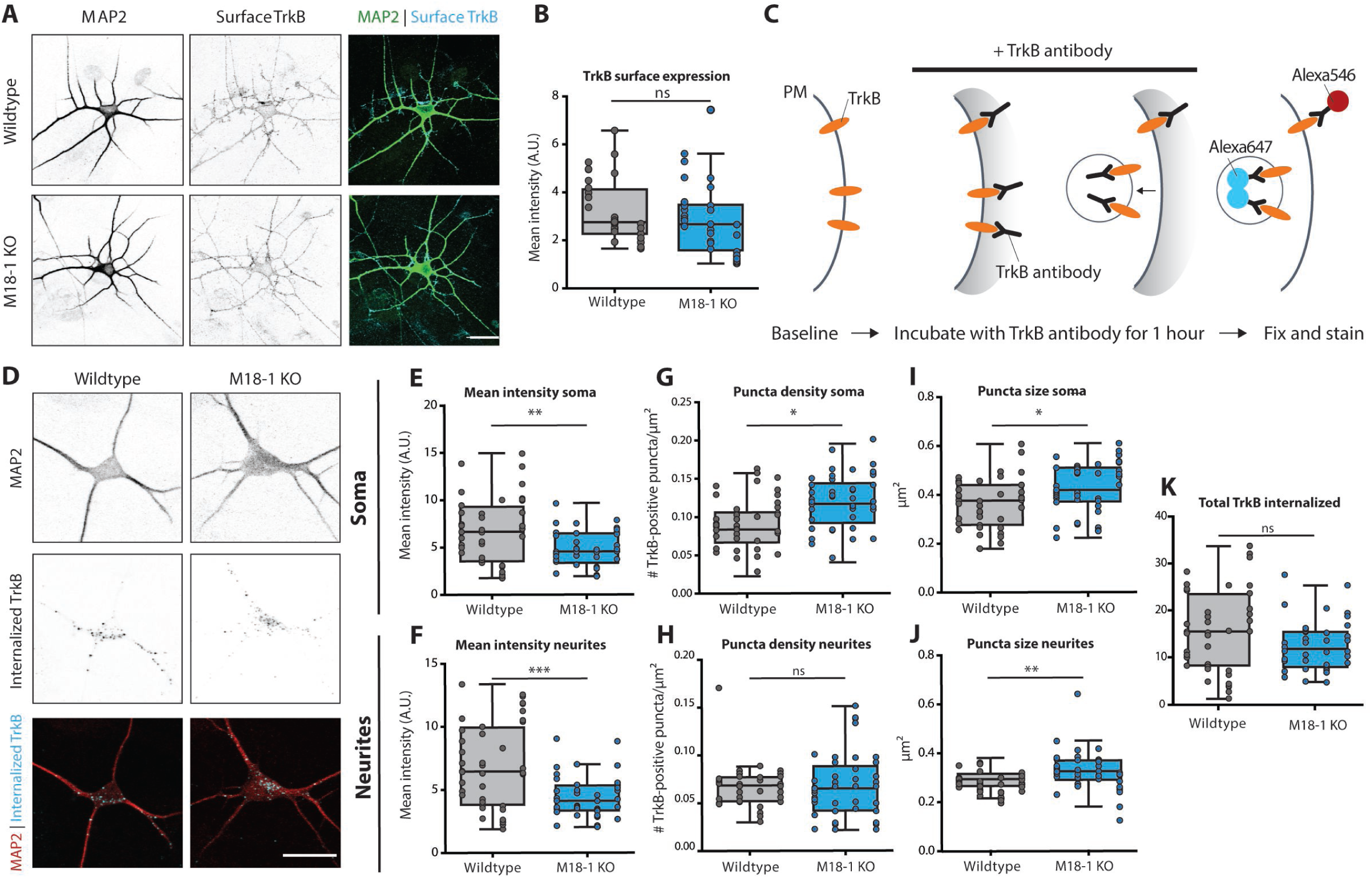
Retrograde TrkB transport is affected in *Munc18-1* KO neurons. A) Typical example of WT and *Munc18-1* KO neurons at DIV3 stained for MAP2 (dendritic marker) and surface TrkB. Scale bar is 25μm. B) Average TrkB surface levels were not affected in *Munc18-1* KO neurons (ns p = 0.23, Mann-Whitney test). C) Cartoon representing TrkB antibody uptake assay. Neurons were incubated with TrkB antibody for one hour, before they were fixed and stained. Remaining surface TrkB was stained with Alexa546, whereas internalized TrkB was stained with Alexa647. D) Typical examples of WT and *Munc18-1* KO neurons fixed after the TrkB antibody uptake assay and stained for MAP2 (dendritic marker) and internalized TrkB (Alexa647). E) Mean TrkB puncta intensity in the soma of *Munc18-1* KO neurons was ∼30% lower (p = 0.003, unpaired T-test). F) Mean TrkB puncta intensity in neurites was ∼35% lower in *Munc18-1* KO neurons (p = 0.0007, Mann-Whitney test). G) *Munc18-1* KO neurons showed an ∼20% increase in TrkB puncta density in the soma (p = 0.01, Mann-Whitney test). H) TrkB puncta density was not different in neurites of *Munc18-1* KO neurons (ns p = 0.12, Mann-Whitney test). I) Average TrkB puncta size in the soma of *Munc18-1* KO neurons was ∼15% bigger (p = 0.01, unpaired T-test). J) In neurites of *Munc18-1* KO neurons, average TrkB puncta size was ∼10% bigger (p = 0.12, Mann-Whitney test). K) Total pool of internalized TrkB was not different between WT and Munc18-1 KO neurons (p=0.07, unpaired T-test). Data is represented in Tukey boxplots. Columns and dots represent individual litters and neurons, respectively.

## Discussion

In the present study we investigated the impact of MUNC18-1 loss in cellular trafficking routes. We show that in absence of MUNC18-1, neurons have a smaller *cis-, medial*-Golgi and TGN, while Golgi stack ultrastructure was normal. ER-to-Golgi and Golgi exit of markers in the constitutive and regulated secretory pathway were unaffected. Retrograde trafficking to the Golgi, however, was impaired.

### *Munc18-1* KO neurons transform from healthy to death within hours

*Munc18-1* KO neurons at DIV3, i.e., during the steepest part of the survival curve, and hours before most neurons have died (Santos et al., 2017), show a strikingly normal morphology when examined at the ultrastructural level. No cellular signs of degeneration as described in previous studies on cell death (e.g., Taatjes et al. 2008) were observed. Hence, *Munc18-1* KO neurons transform from morphologically normal, healthy neurons to cell death within hours, with only a smaller Golgi/TGN as a morphological indicator of the nearing cell death. In addition, reduced expression of several, but not all proteins was detected at this stage (Fig. 4 & 5), which is a common feature in other cell death models (e.g., Patel et al. 2002). Cell death in other cell types is known to also occur within hours, e.g. at a rate of 5% per hour (Wolbers et al., 2004), but such high rates are induced by lethal cytotoxic compounds or radiation (Jessel et al., 2002; Wolbers et al., 2004; Forcina et al., 2017). In conclusion, the transformation from normal, healthy neurons to death is remarkably fast in *Munc18-1* KO neurons. The cell autonomous defects caused by MUNC18-1 loss, and probably also syntaxin-1 or SNAP25 loss, trigger cell death with exceptionally fast kinetics, comparable to cytotoxic poisoning.

### MUNC18-1 deficiency leads to atypical condensation in all aspects of the Golgi

The earliest phenotype detected upon loss of MUNC18-1 is a smaller and rounder Golgi area, suggesting condensation of the Golgi complex. However, Golgi stacking and ultrastructure cisternae morphology were not different. In addition, the average intensity of a Golgi-resident marker was not increased in *Munc18-1* KO neurons. This is strikingly different from classical Golgi condensation phenotypes, where the round Golgi morphology is accompanied by swollen cisternae and/or accumulation of Golgi resident markers (Bard et al., 2003; Young et al., 2005; Lázaro-Diéguez et al., 2006; Dippold et al., 2009). The clear absence of ultrastructure abnormalities places MUNC18-1-induced Golgi abnormalities in a novel category, distinct from typical Golgi condensation associated with cell death.

### MUNC18-1’s role in actin regulation does not explain Golgi abnormalities

The characteristic elongated morphology of the Golgi is maintained by a complex interplay between lipids and the actin cytoskeleton (Dippold et al., 2009). MUNC18-1 has an established role in controlling actin organization via a hydrophobic residue in β-sheet 10 (Pons-Vizcarra et al., 2019). We showed that depolymerisation of actin in WT neurons by LatB treatment phenocopied the smaller and rounder Golgi area observed upon MUNC18-1 loss, indicating that actin defects might explain the observed Golgi abnormalities. However, Golgi morphology was restored upon expression of a MUNC18-1 mutant that fails to support a normal actin organization. Moreover, previous studies investigating Golgi abnormalities upon actin depolymerisation reported perforation/fragmentation and severe swelling of Golgi cisternae (Lázaro-Diéguez et al., 2006), and defects in VSVG transport (Dippold et al., 2009). This is in contrast with the observations in *Munc18-1* KO neurons in the present study; cisternae morphology was normal and VSVG transport was not affected. Hence, it is unlikely that defects in the actin cytoskeleton explain the Golgi abnormalities in *Munc18-1* KO neurons.

### Anterograde/retrograde trafficking imbalance explains Golgi abnormalities

Golgi morphology and function are tightly linked. A previously published RNAi screen showed that 75% of the proteins that regulate Golgi morphology, also played essential roles in Golgi function (Chia et al., 2012). Since loss of MUNC18-1 results in a severe reduction of Golgi membrane, it was anticipated that Golgi function would be disturbed. Remarkably, anterograde trafficking or Golgi exit were normal for multiple cargo markers in the constitutive and regulated secretory pathway. In addition, it is unlikely that glycosylation, one of the main functions of the Golgi, is affected since glycosylation of TrkB is essential for its correct localisation in the cell (Watson et al., 1999; Mutoh et al., 2000). Hence, Golgi-intrinsic functions and Golgi-export are not affected by MUNC18-1 deficiency.

In contrast, two independent assays uncovered retrograde trafficking defects in *Munc18-1* KO neurons. First, retrograde transport of CTB from the PM to the Golgi was reduced in *Munc18-1* KO neurons. Second, retrograde TrkB trafficking was disturbed. Upon endocytosis, CTB and TrkB are both sorted to the endocytic pathway in Rab7-positive organelles and transported back to the soma (Ehlers et al., 1995; Bhattacharyya et al., 1997). These organelles, referred to as signalling endosomes, engage interactors to enable downstream signalling pathways (Delcroix et al., 2003; Villarroel-Campos et al., 2018). Once in the soma, signalling endosomes are either directed to lysosomes, or fuse with the TGN to recycle proteins and lipids (Progida and Bakke, 2016; Villarroel-Campos et al., 2018). For both assays, it is unlikely that the defects are explained by impaired endocytosis or receptor recycling: for CTB retrograde transport to the Golgi was impaired after normalizing for the total internalized CTB pool (Fig. 6K), and in the TrkB assay the total pool of internalized TrkB was unaffected (Fig. 7K). Hence, we conclude that the retrograde trafficking defects occur after the initial endocytosis of CTB/TrkB. Indeed, perturbations in retrograde transport of signalling endosomes in Endophilin triple KO neurons or inhibition of dynein motor proteins result in similar cellular phenotypes and reduced survival, albeit less pronounced, as presented in this study (Burk et al., 2017; Budzinska et al., 2020). Hence, defects in retrograde transport of signalling endosomes provide a plausible explanation for the observed abnormalities in CTB transport and retrograde TrkB trafficking, and also for the reduced Golgi size in *Munc18-1* KO neurons. A proper balance between the anterograde and retrograde pathways is essential to maintain Golgi organization and architecture (Pavelka and Ellinger, 2008). Previous studies in non-neuronal cells have shown that the disruption in retrograde endosome-to-Golgi transport, but not anterograde transport, results in abnormal Golgi morphology, albeit with distinct morphological hallmarks than observed in the current study (Naslavsky et al., 2009; Shitara et al., 2013). Hence, the imbalance between antero- and retrograde pathways provides an explanation for the observed Golgi abnormalities in *Munc18-1* KO neurons.

### A potential role for Munc18-1 and its t-SNAREs in SNARE-dependent fusion between endosomes and Golgi

Together, our data suggest a role for MUNC18-1 in neuronal endosome-to-TGN transport, resulting in abnormalities in Golgi morphology. Previously, we have shown that similar Golgi abnormalities occur in the absence of its t-SNARE partners (Santos et al., 2017). These proteins are known to work together in membrane fusion reactions (Sudhof, 2013) and a defective fusion reaction is therefore the most plausible explanation for the common Golgi phenotypes and neurodegeneration in the absence of these three proteins. Endosome-to-TGN retrograde trafficking relies on multiple SNARE-dependent fusion reactions (Mallard et al., 2002). Interfering with these fusion reactions causes abnormalities in Golgi morphology, while cisternae ultrastructure and anterograde transport remained unaffected (Rahajeng et al., 2010; Shitara et al., 2013), i.e., similar to the situation described in the current study for MUNC18-1 deficiency. These apparent similarities suggest that MUNC18-1 and its cognate tSNAREs also play a role in endosome-to-TGN SNARE-dependent fusion. However, in non-neuronal cells, dedicated S/M proteins have been described to operate in such retrograde pathways, especially mVps45 (Rahajeng et al., 2010) and possibly also mSly1 (Laufman et al., 2009), and to organize SNARE-dependent fusion with specialized Q-SNAREs syntaxin-6, and −16 (Mallard et al., 2002), and probably also syntaxin-5 (Tai et al., 2004; Amessou et al., 2007) and syntaxin-10 in other mammals (see for reviews, (Toonen and Verhage, 2003; Dingjan et al., 2018)). In neurons, little is known about the function of these S/M proteins. MUNC18-1 and syntaxin-1 may account for a distinct, neuron specific, retrograde pathway, and/or their loss may indirectly affect the equilibrium in cellular levels or availability of other SNAREs and SNARE-organisers in retrograde cellular trafficking, causing the observed retrograde transport and Golgi abnormalities.

### Defects in retrograde pathways can explain degeneration in *Munc18-1* KO brains

The impairment in retrograde transport might not only provide an explanation for the Golgi abnormalities, but also for the fact that *Munc18-1* KO neurons die. Correct retrograde transport of TrkB is essential for neuronal survival (Reichardt, 2006; Burk et al., 2017), and comparable abnormalities in TrkB-containing endosomes have been reported upon perturbations of the retrograde pathways, leading to defective neurotrophic signalling and reduced neuronal survival (Wan et al., 2008; Budzinska et al., 2020). In addition, enlarged signalling endosomes have directly been linked to neurodegenerative processes (Xu et al., 2016). It was previously shown that administration of BDNF to *Munc18-1* KO neurons delays the degeneration (Heeroma et al., 2004), strengthening the notion that neurotrophic signalling is involved in MUNC18-1-dependent viability. Together, this suggests that defective retrograde pathways in *Munc18-1* KO neurons result in several intracellular impairments, including neurotrophic signalling, ultimately leading to cell death.

## Acknowledgments

This work was supported by the European Union ERC Advanced Grant 322966 to M.V. and Horizon 2020 grant COSYN (RIA grant agreement n° 610307, to M.V.) and by NWO Gravitation program BRAINSCAPES (NWO: 024.004.012). We thank Desiree Schut and Lisa Laan for preparing glia feeders and culturing neurons; Robbert Zalm for virus production and cloning; Joost Hoetjes and Joke Wortel for breeding and genotyping mutant mice.

## Notes

**Conflict of interest statement:** The authors declare no competing financial interests.

